# High-throughput, image-based screening of genetic variant libraries

**DOI:** 10.1101/143966

**Authors:** George Emanuel, Jeffrey R. Moffitt, Xiaowei Zhuang

**Affiliations:** Howard Hughes Medical Institute, Cambridge, MA 02138, USA; Graduate Program in Biophysics, Cambridge, MA 02138, USA; Department of Chemistry and Chemical Biology, Cambridge, MA 02138, USA; Department of Physics, Harvard University, Cambridge, MA 02138, USA

## Abstract

Image-based, high-throughput, high-content screening of pooled libraries of genetic perturbations will greatly advance our understanding biological systems and facilitate many biotechnology applications. Here we introduce a high-throughput screening method that allows highly diverse genotypes and the corresponding phenotypes to be imaged in numerous individual cells. To facilitate genotyping by imaging, barcoded genetic variants are introduced into the cells, each cell carrying a single genetic variant connected to a unique, nucleic-acid barcode. To identify the genotype-phenotype correspondence, we perform live-cell imaging to determine the phenotype of each cell, and massively multiplexed FISH imaging to measure the barcode expressed in the same cell. We demonstrated the utility of this approach by screening for brighter and more photostable variants of the fluorescent protein YFAST. We imaged 20 million cells expressing ~60,000 YFAST mutants and identified novel YFAST variants that are substantially brighter and/or more photostable than the wild-type protein.

---

High-throughput screening of genetic perturbations is playing an increasingly important role in advancing biology and biotechnology. For example, by observing the effects of a large number of amino acid changes within a selected protein, large-scale screening can enable efficient searches for fluorescent proteins better adapted for bioimaging or protein and nucleic acid drugs with desired therapeutic properties. It can also enable the examination of how mutations of a protein affect cell function or physiology. Since each cell is composed of many genes, large-throughput screening can also allow the effects of inhibition or activation of individual genes or combinations of genes to be tested at the genomic scale, which will help decipher the effects of genes and gene networks on cellular behaviors.

Large-scale screening efforts are greatly facilitated by pooled, high-diversity libraries of genetic perturbations because of the ease and scalability associated with the construction of these pools. By using methods such as error-prone PCR^1^ or cloning with large, defined pools of array-synthesized oligonucleotides^2^, it is often possible to create pooled libraries with a very large number of genetic variants or perturbations using a similar degree of effort as required to make any individual library member. The screening of pooled libraries then depends critically on the ability to measure not only the desired phenotype but also the genotype of the corresponding library members. The measured phenotypes can be simple in some cases, such as protein affinity as measured by standard pull-down assays^3,4^, or cellular fluorescence as determined by FACS^5^, and in both cases, the genotype can be determined via sorting or enriching library members with the desired phenotype followed by techniques such as sequencing to determine the genotype. However, there are many phenotypes that cannot be quantified with these techniques. Phenotypes ranging from cellular morphology and dynamics to the intracellular organization of different cellular components require high-resolution, time-lapse optical microscopy to be measured. Unfortunately, it has proven challenging to combine pooled library screening with high-resolution optical microscopy because it is difficult to isolate or recover individual library members based on their imaged phenotype and then determine their genotype. If, however, one had the ability to measure the genotype of individual library members also by imaging, then library member isolation and recovery would not be required, making it possible to combine the ease and scalability of pooled library screening with imaging-based phenotype measurements.

Here we report a novel high-throughput, imaging-based screening method that allows the characterization of both phenotype and genotype for pooled populations of genetically diverse cells. In this method, we associate each genetic variant with a unique barcode composed of a series of short oligonucleotide hybridization sites. After introducing the barcoded genetic variant library into a population of cells and measuring phenotypes with imaging, the cells are fixed and the barcodes are determined using multiplexed error robust fluorescence in situ hybridization (MERFISH), a method that utilizes combinatorial labeling and sequential imaging to identify a large number of barcodes^6^. We tested the feasibility and quantified the accuracy of this screening approach by screening a library of 1.5 million cells with 80,000 unique barcodes in which cells either did or did not express the fluorescent protein mMaple3^7^. Based on these measurements, we estimated that our genotype misidentification rate is less than 1% for individual cells. To demonstrate the power of this approach, we utilized it to improve the brightness and photostability of YFAST, a recently developed fluorescent protein that becomes fluorescent upon binding to an exogenous chromophore^8^. With this approach, we efficiently screened 20 million *E. coli* cells containing ~160,000 unique barcodes and ~60,000 unique YFAST mutants, which resulted in the identification of YFAST variant with substantially increased brightness and photostability. By utilizing high-resolution imaging, we envision that this approach also has the potential to screen the effect of genetic perturbations on other cellular properties that would be difficult to measure with non-imaging methods, such as the morphology and dynamics of the cell, or the intracellular distributions of proteins/RNAs and the spatial organization of the genome.

In our previous demonstration of MERFISH^6^, we utilized this approach to massively increase the multiplexing of single-molecule fluorescence in situ hybridization^9,10^ and measure a large number of distinct RNA species simultaneously in single cells^6^. We subsequently improved the throughput of MERFISH to allow a large number of cells to be measured in a short period of time^11^. Here we reasoned that we could use this approach to identify the genotype of individual cells if each cell contains a unique DNA barcode that is associated with the gene variant of interest. Thus, cells could be identified by effectively measuring the identity of the RNA that it expressed from the barcode. To construct such barcodes, we designed nucleotide sequences that contain a concatenation of a fixed number of hybridization sites, where the sequence of each hybridization site was one of a pair of unique sequences associated with that site (Fig. 1A). We term these sequences readout sequences. Effectively each barcode sequence can be thought of as representing an N-bit binary code, where there is a unique readout sequence to represent a “1” or a “0” in each bit. Thus, each barcode is comprised of a set of *N* hybridization sites drawn from 2N unique readout sequences: readout sequence 1-0, readout sequence 1-1, readout sequence 2-0, readout sequence 2-1, …, readout sequence *N*-0, readout sequence *N*-1. For example, a barcode sequence corresponding to the binary word 101…1 would consist of readout sequence 1-1, followed by readout sequence 2-0, then readout sequence 3-1, …, and finally readout sequence *N*-1. A similar approach could be used to construct alternative barcodes using more unique readout sequences at each site, i.e. three sites would correspond to a ternary code, or using the absence of a site as a measurable signal, i.e. a ‘1’ could be the presence of a site while a ‘0’ could be represented by its absence.

**Fig. 1.**
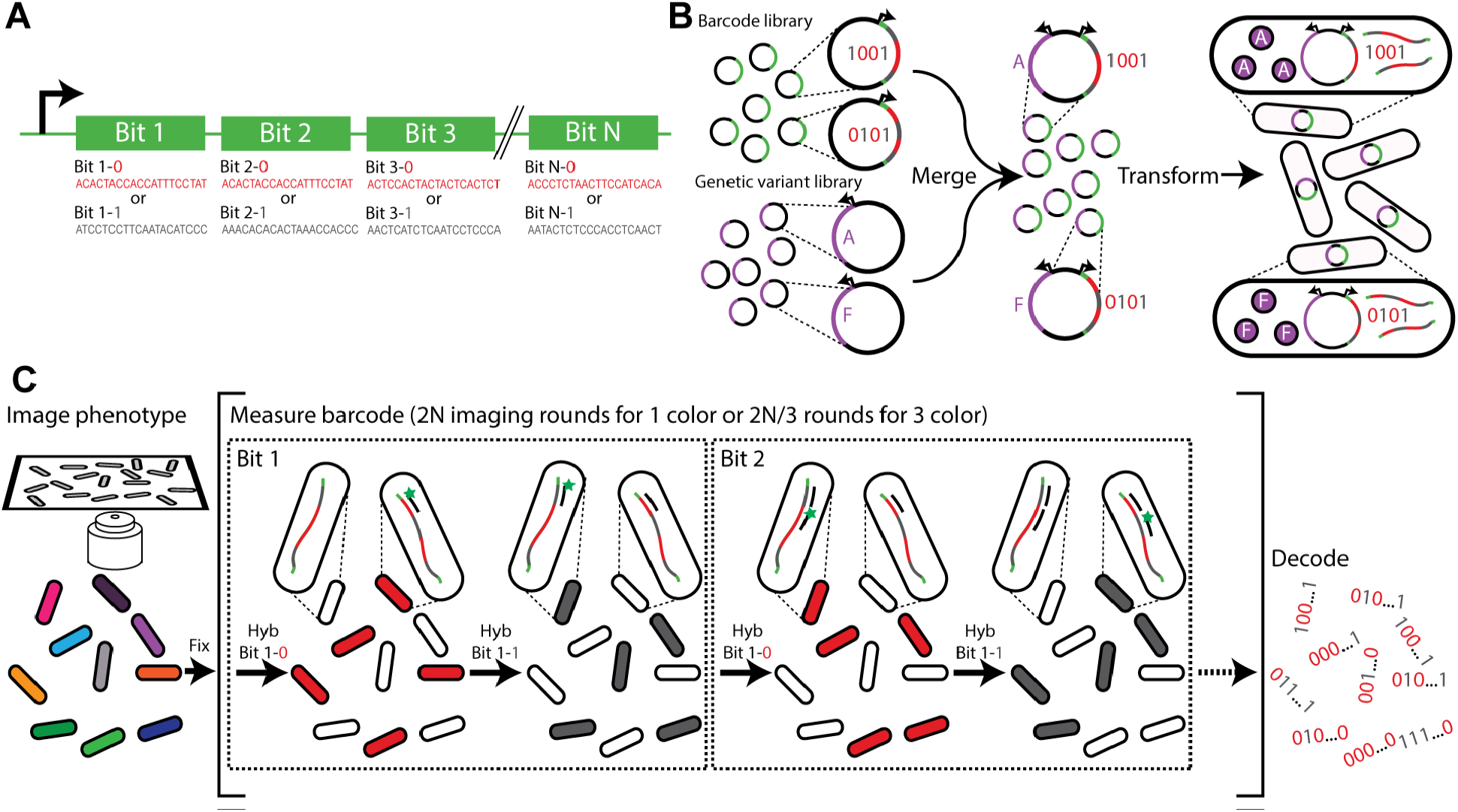
A high-through, image-based screening method using massively multiplexed fluorescence in situ hybridization. (A) Schematic depiction of an RNA barcode. Each RNA barcode consists of the concatenation of a series of hybridization sequences, each of which can be associated with a different bit in an *N*-bit binary barcode. Each hybridization sequence can utilize one of two readout sequences unique to that position, with one readout sequence associated with a “1” at that bit and another with a “0”. (B) Schematic depiction of library construction. The library of barcodes is merged with a library of genetic variants and transformed into bacteria. The correspondence between the barcodes and genetic variants is determined by sequencing. (C) Schematic diagram of the image-based phenotype-genotype characterization. The phenotype is first characterized in surface-adhered cells. Then, the cells are fixed, and multiple rounds of hybridization are used to measure the barcodes. During the first round, readout probe 1-0 is added and cells with barcodes that read “0” in the first bit, i.e. which contain the readout sequence 1-0, should bind to the probe and become fluorescent, whereas cells with barcodes that read “1” in the first bit should remain dark. Once readout probe 1-0 is extinguished, readout probe 1-1 is added and the cells with barcodes that read “1” in the first bit, which contain the readout sequence 1-1, should become fluorescent. This difference in fluorescence intensity allows the value of bit 1 to be determined for each cell. This process is repeated for the remaining bits. After measuring all bits, the barcode is determined, revealing the identity of the genetic variant contained in the cell and linking the measured phenotype to the genotype.

As an illustration of one way in which this barcoding scheme could be used to identify genetic variants, we created a library of plasmids expressing these N-bit barcodes and a library of plasmids expressing a series of genetic variants of a protein of interest. To create a barcoded library of genetic variants, we fused the barcode sequences and the sequences expressing the genetic variants to create a library of new plasmids, each of which expresses a random combination of a genetic variant and a barcode, and introduced the library into *E. coli* cells such that each cell only expresses one plasmid (Fig. 1B) (see Materials and Methods for the details). To reduce the chance of a barcode appearing in the library associated with more than one genetic variant, we bottlenecked the diversity of the barcoded genetic variant library by limiting the number of cells expressing the barcoded genetic variant library to 1 to 10% of the total barcode diversity of 2^*N*^. We then determined which barcode is associated with which genetic variant by extracting these plasmids and sequencing them with next generation sequencing to construct a look-up table. These measurements also detected a remaining small fraction of barcodes that were each associated with more than one genetic variant. If detected, these barcodes were removed from further analysis.

To screen this barcoded genetic variant library, we imaged cellular phenotypes while the cells were still alive (Fig. 1C). Then, the cells were fixed without removing them from the microscope, and the RNA expressed from the barcodes were read out using multiplexed error-robust fluorescence in situ hybridization (MERFISH)^6^.

During the barcode readout process, multiple hybridization rounds were used and fluorescently labeled readout probes complementary to each readout sequence on the barcode were hybridized in each round to detect which readout sequences are present in which cells. First, readout probe 1-0, complementary to readout sequence 1-0, was introduced so that it can hybridize to cells that contain readout sequence 1-0, namely the cells containing barcodes whose first bit is “1”, causing those cells to become brightly fluorescent. All the cells were imaged and then the fluorescence signal removed from the sample. Then, readout probe 1-1 was hybridized. Since every barcode contains either readout sequence 1-0 or readout sequence 1-1, the cells that did not become fluorescent in the first round should became fluorescent in the second round. The value for bit 1 for each cell was then assigned based on the fluorescence intensity ratio between these two rounds. This process was iterated until all *N* bits were probed. To reduce the number of hybridization rounds, we used multiple color imaging with spectrally distinct fluorescent dyes to probe for the presence of multiple readout sequences simultaneously in each round^11^. Three-color imaging was used in this work though a readout scheme using more colors is also possible.

Since each cell expresses many copies of its corresponding barcode RNA, the fluorescence signal was very bright, and hence the readout error rate for each bit was very small. Thus, we did not find it necessary to use error correcting codes in this case, as we previously used for MERFISH^6^, but instead all 2^*N*^ possible barcodes could be used in principle. However, in practice, to avoid a barcode appearing paired with multiple mutants in the same library, we bottlenecked the number of unique library members, as described above. The use of only a subset of barcodes allowed the experimental measurement of the frequency with which unused barcodes were detected, which in turn allowed us to quantify the rate of misidentifying the genotype of a cell, as we describe below. With a constant degree of bottlenecking, this type of internal error measurement and error robustness is maintained even as the number of possible binary barcodes increases exponentially with the number of bits. Thus, this barcoding scheme should allow millions of unique barcodes to be measured with only tens of hybridization rounds.

To test the accuracy of this screening approach, we created a library containing only two “genetic variants”, the mTagBFP2 gene and the fusion of mTagBFP2 and mMaple3 genes (Fig. 2A). Two barcoded libraries were created by merging a 21-bit binary barcode library, consisting of more than 2 million (2^21^) unique barcodes, with the two plasmids (one containing the mTagBFP2 and the other containing mTagBFP2-mMaple3 gene), one for each library, by isothermal assembly. Then, each of these complete libraries was electroporated into *E. coli* cells, and a fixed number of cells were extracted to bottleneck these libraries to ~40,000 unique members. The plasmids were extracted from the cells and sequenced to determine which of the ~2 million possible barcodes were present in each library and, thus, associated with the presence or absence of mMaple3. Sequencing confirmed that in the mixture of the two libraries, ~80,000 unique barcodes were present, as expected, representing 4% of all 2^21^ possible barcodes.

**Fig. 2.**
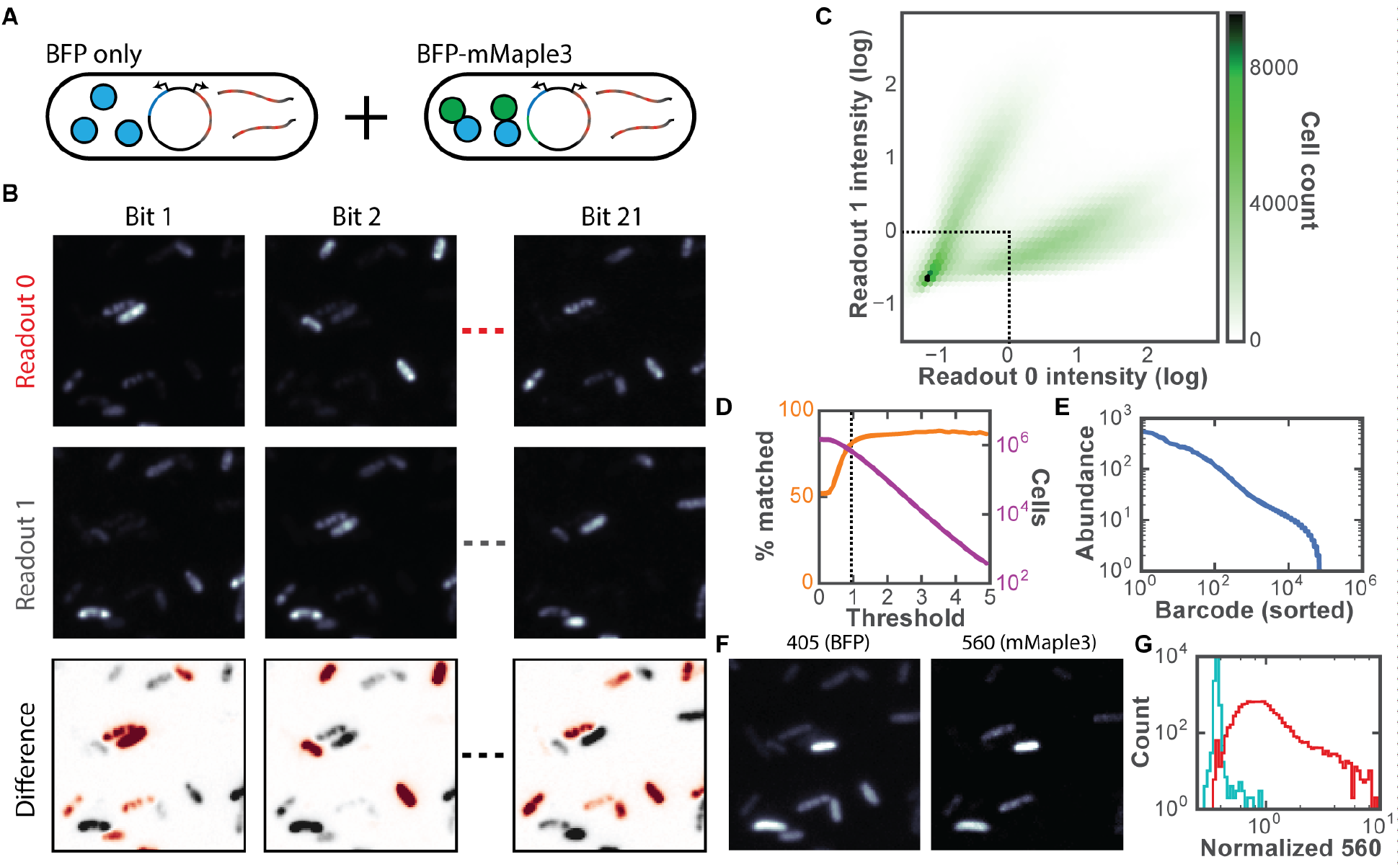
Performance characterization of the screening method by measuring 600,000 cells containing 80,000 unique barcodes associated with two known genotypes and phenotypes. (A) Schematic diagram of the library constituents. Among the 80,000 distinct 21-bit barcodes, half are associated with the mTagBFP2 gene while the other half are associated with the mTagBFP2-mMaple3 fusion gene. Both sets of cells will express RNA barcodes, but only those expressing mTagBFP2-mMaple3 will be fluorescent in the 560-nm channel. (B) Fluorescent images for each readout from a subset of the full 21 bits. Images after the readout sequence corresponding to a “0” (top) or to a “1” (middle) are hybridized are shown. The difference image (bottom) indicates whether a “0” or “1” is assigned to the barcode within that cell in that bit (red and gray indicate “0” and “1”, respectively). (C) Two-dimensional histogram of normalized fluorescence intensities for readout 0 and readout 1 of bit 1 for each cell. The fluorescence intensities are normalized to the median values. The dotted line depicts the threshold used for eliminating cells that appear dim in both readouts. The shade of green indicates the number of cells. (D) The percent of barcodes decoded in the imaging experiment that match barcodes determined to be in the library by sequencing (orange) and the number of cells (magenta) above the readout intensity threshold with varying threshold magnitude. The dotted line corresponds with the threshold of 1 shown in (C). (E) Abundance of each barcode. The abundance is the number of cells in the imaging experiment assigned to each barcode. (F) Fluorescence image of mTagBFP2 and fluorescence image of post-activation mMaple3 of the same region as (B). (G) Histograms of median mMaple3 fluorescence intensity normalized to mTagBFP intensity for barcodes associated with the mMaple3-mTagBFP2 fusion gene (red) and for those associated with the mTagBFP2-gene (cyan).

We then characterized this combined library using our screening strategy. We first measured the fluorescence properties of the cells expressing mTagBFP2 or mTagBFP2-mMaple3 by illuminating with 405-nm light to measure mTagBFP2 fluorescence, illuminating with 405-nm light for an additional ~4s in order to switch the mMaple3 protein to its red-shifted fluorescent state, and then measuring the fluorescence intensity of the red-shifted mMaple3 by illuminating with 560-nm light. We then fixed the cells with methanol and acetone and read out the barcodes in each cell using the procedure described above. We utilized alcohol fixation as opposed to crosslinking fixatives such a paraformaldehyde since it has been established that alcohol fixation greatly enhances the rate of hybridization to RNA^12^.

During the barcode reading step, we examined the fluorescence signal observed for individual cells during different rounds of hybridization and imaging. Indeed, as expected, we found that cells that were bright for one readout of a given bit were dim for the other readout (Fig. 2B). For all 1.5 million cells observed, a two-dimensional (2D) histogram of the bit 1 measurements, i.e. the fluorescence intensities determined in the probe 1-0 imaging round and probe 1-1 imaging round, was constructed, and this histogram suggests there are two distinct populations of cells (Fig. 2C). The first population appeared bright when hybridized to probe 1-0 and dim when hybridized to probe 1-1 while the second population appeared dim when hybridized to probe 1-0 and bright when hybridized to probe 1-1. This observation is consistent with the readout sequence 1-0 being present in the first population and the readout sequence 1-1 being present in the second population. However, a substantial fraction of cells appeared dark in both imaging rounds, possibly because they are not expressing sufficient barcode RNA, or they are insufficiently permeabilized for readout probe hybridization. We therefore used a thresholding strategy to remove these dim cells from further analysis. Specifically, we require that the “0” or “1”readout signal for each bit is larger than the median intensity observed for that readout signal across all cells. More than 600,000 measured cells satisfied this conservative criterion, and the barcodes expressed in these cells were determined. Among these cells, 84% of the measured barcodes matched a barcode contained in the library as determined by sequencing (Fig. 2D).

For the unmatched 16% of the cells, an experimental error must have occurred. Either the barcode was present in the library but it was not detected by sequencing or the barcode was not present in the library and an error occurred during barcode imaging that misidentified the barcode. While we did not use these cells in further analysis, the presence of this unmatched fraction can be used to estimate our experimental error rates in barcode identification. We first note that the library only contains 4% of all possible barcodes for the 21-bit binary encoding used, as described above, so assuming that a readout error in the imaging process is equally likely to result in a cell being assigned any of the 2^21^ barcodes, there is a 96% chance that the error results in identifying a barcode that does not match one in the library (type 1 error). Next, we denote the probability that the barcode in a cell is incorrectly determined as x. Then this probability x multiplied by 96% should be equal to 16%, the fraction of cells that we found containing barcodes that did not match anyone in the library. Hence, x should be equal to 0.167 and the probability that the error results in a different barcode that is present in the library should be only 4% of x, which is ~0.67% (type 2 error). We note that only type 2 errors would affect our final results because only these errors have the capacity to generate a misidentified genotype. Type 1 errors would be detected as unmatched barcodes and discarded. Therefore, our estimated misidentification rate, i.e. the probability that both an error occurs and the error yields a barcode this is already present in the library, is less than one percent.

The fidelity of the barcode measurement can also be verified by comparing the measured phenotype of cells from the mMaple3 fluorescence level to the phenotype predicted by the measured barcode. We used the phenotype measurements described above to determine the mTagBFP2 fluorescence intensity and the mMaple3 fluorescence intensity of each cell (Fig. 2F). All cells that contained the same measured RNA barcode were grouped and the median ratio of mMaple3 intensity to mTagBFP2 intensity was calculated for each group containing at least 5 cells. Since from sequencing we know which barcodes should be associate with mMaple3, we calculated the median intensity ratio for cells identified for each of these barcodes and constructed a histogram of these ratios. Using the same approach, we constructed a histogram of the ratios for the barcodes known to only be associated with mTagBFP2 (Fig. 2G). These two distributions are largely separated with only a small overlap. We set a threshold based on the intersection point of the two histograms such that the cells with fluorescence intensity ratios larger than this threshold were classified as containing the mTagBFP2-mMaple3 fusion protein and the cells with the intensity ratio below the threshold as containing mTagBFP2. We then compared these phenotype assignments based on fluorescence intensity to the genotypes based on the measured barcodes. We found that less than 1% of cells have a phenotype and genotype disagreement, indicating a rate of misidentification comparable to that estimated above. However, we note that this error rate is likely an overestimate since the natural spread in the intensity distribution of cells in each group should allow from some overlap of these distributions. Based on the above quantifications, we conclude that our high-throughput imaging-based screening approach is capable of accurately associating the phenotype with the genotype in a large number of cells.

To demonstrate the utility of our approach for screening a large library of mutants to find proteins with desired properties, we screened for improved variants of a recently developed fluorescent protein, YFAST. YFAST is not itself fluorescent but only becomes fluorescent upon binding to an exogenous, GFP-like chromophore, such as HBR or HMBR (Fig. 3A)^8^. We created a library of YFAST variants, merged it with our 21-bit barcode library, and sequenced the resulting plasmid library to build the lookup table between variant and barcode.

**Fig. 3.**
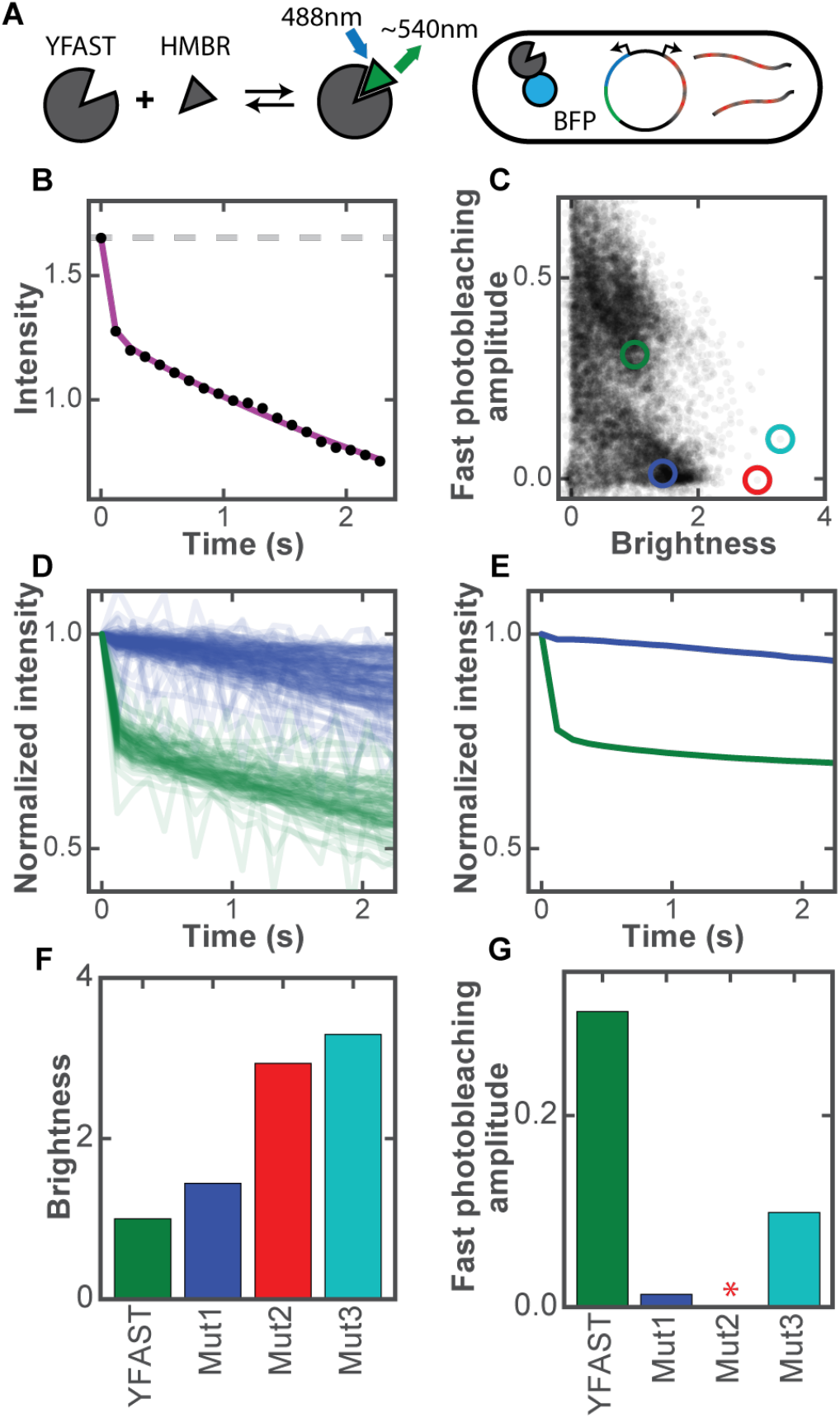
Screening YFAST mutant libraries for improved brightness and photobleaching kinetics. (A) Schematic diagram of YFAST library design. YFAST is dark on its own, but it becomes fluorescent upon binding to the ligand, for example HMBR^8^. A library of YFAST variants fused to mTagBFP2 for normalization is merged with a library of barcodes and transformed into *E. coli* cells. (B) The initial intensity and the photobleaching curve (black circles) of the original YFAST measured from a single cell in the library screen measurement. The initial intensity (gray dashed line) is measured as the intensity of YFAST fluorescence under 488-nm illumination after background subtraction normalized to the mTagBFP2 fluorescence intensity under 405-nm illumination. The photobleaching curve (circles) is measured by illumination with 488-nm light to excite YFAST only. The curve is fit to a double exponential decay (red line) with the background level determined by the intensity of cells that have dark YFAST variants. (C) Scatter plot of the relative brightness and fast-photobleaching amplitude for each measured mutant. Each point depicts the median brightness and fast-photobleaching amplitude of all cells associated with one mutant in the library. The brightness values are normalized to that of the original YFAST. Here only the mutants that contain at least 10 imaged cells are depicted. The original YFAST (green) and several selected mutants (blue, red, and cyan) with greater brightness and/or much smaller fast-photobleaching amplitudes are marked. (D) The photobleaching curves for individual cell corresponding to the original YFAST (green) and one selected mutant (blue dot in (C)) from the library measurement. The fluorescence intensities of the initial time point are normalized to 1. (E) The average bleaching curves for the original YFAST (green) and one selected mutant (blue dot in (C)) measured in isolation in pure culture. The original YFAST and mutant were individually expressed in the *E. coli* cells together with a mTagBFP2 using the same plasmid construct as described for the library constructs. The initial brightness values of YFAST and mutant are normalized as described in (D). (F and G) Bar charts of the relative brightness values (F) and fast-photobleaching amplitudes (G) for several selected mutants as marked in (C) by the same colors. The brightness values are normalized to that of the original YFAST. * indicates a value close to zero and hence not visible in the bar graph.

We sought to simultaneously improve two properties of YFAST: the fluorophore brightness and the photobleaching kinetics. We note that while fluorophore brightness is a property that can be measured and selected via more traditional screening methods, e.g. FACS, photobleaching kinetics require a time-lapse measurement of the fluorescence from a single cell and, thus, would be challenging to perform with other approaches. In particular, we observed that the photobleaching decay of YFAST follows a double exponential decay with one component decaying much faster than the other. We sought to identify mutants that eliminate this fast decaying component.

To measure the brightness of different YFAST variants while controlling for potential variations in the expression level, we fused YFAST variants to mTagBFP2 and imaged individual cells with both 405-nm and 488-nm illumination, respectively (Fig. 3A). The relative brightness of YFAST was calculated as the ratio of the background subtracted YFAST fluorescence intensity measured with 488-nm illumination in the presence of the chromophore to the mTagBFP2 intensity measured with 405-nm illumination in the absence of the chromophore. To characterize the photobleaching kinetics of YFAST variants, we measured the decrease in intensity over 20 frames under constant 488-nm illumination alone (Fig. 3B).

We constructed a series of YFAST variant libraries that contain (1) all possible single amino acid substitutions, insertions, and deletions (termed single-amino-acid libraries), (2) multiple mutations surrounding the chromophore binding pocket in each library member (chromophore adjacent library), (3) a combination of the best mutations identified from the chromophore adjacent library and the best mutations identified from the single amino acid libraries that are also near the chromophore, or (4) all possible single amino acid substitutions, insertions, and deletions based on a favorable mutant derived from the above libraries. In total, we constructed libraries containing roughly 60,000 unique YFAST variants associated with ~160,000 barcodes, and we used our high-throughput, image-based screening method to measure the brightness, photobleaching kinetics, and genotype for this library of variants across 20 million total cells. From each cell’s photobeaching decay curve, we determine the relative amplitude of the fast decay component (fast-photobleaching amplitude). We then grouped cells based on the genotypes measured and computed the median brightness and fast-photobleaching amplitude for each of these cell groups (Fig. 3C).

We observe a wide range of relative brightness values and fast-photobleaching amplitudes. To confirm that these variations represent true phenotypic variability, we first selected two variants, the wildtype YFAST (green dot in Fig. 3C) and a variant with a much smaller (nearly eliminated) fast-photobeaching amplitude (blue dot in Fig. 3C), and plotted the measured photobeaching decay curves for the hundreds of measured individual cells that contained the two barcodes associated with these genotypes. The photobleaching decay curves measured for these two sets of cells clearly separate into two populations that correlate strongly with their genotypes, indicating a high degree of reproducibility in the measurements of phenotypes within individual cells. Next, we individually cloned these two YFAST variants and measured their properties in pure culture. We observe nearly identical improvement in photobleaching kinetics in these measurements as we did when these phenotypes were measured in the context of the variant library. By screening ~60,000 variants of YFAST, we identified mutants that are substantially brighter, or mutants that have eliminated the fast-photobeaching component, or mutants with improvements in both aspects (Fig. 3F, G). Moreover, because we have an exact genotype measured for each phenotype, we have produced a rich dataset of YFAST mutations and their phenotypic consequences that could be examined to extract information on both the biophysical properties of this protein as well as information that could guide future mutational screens.

In summary, we developed a method for image-based screening of large genetic variant libraries by coexpressing the genetic variants and barcode that can identify these genetic variants in cells, and determining both the phenotypes of the genetic variants and the barcodes in the same cells using imaging. By reading out barcodes using massively multiplexed FISH, we demonstrated the ability to screen hundreds of thousands of barcodes that correspond to tens of thousands of unique genetic variations. Using this approach, we identified mutations in the YFAST protein, a recently discovered ligand-dependent fluorescent protein, with substantially improved brightness and photostability. We expect that this novel high-throughput, image-based screening method can be applied broadly to improving properties or identifying new properties of proteins and nucleic acids, as well as to deciphering the effects of genes and gene networks on cellular/organism behaviors at the genomic scale.

## Materials and Methods

### Barcode library assembly

The barcode library consists of a set of plasmids, each containing a DNA barcode sequence that encodes a RNA designed to represent a single 22-bit binary word that is transcribed by the lpp promoter. Every barcode in the library has exactly 22 readout sequences, one corresponding to each bit, designed to be read out by hybridizing fluorescent probes with the complementary sequence. Although 22 bits are present in the barcode, to reduce the number of hybridization rounds, experiments were conducted reading out either 18 or 21 of the possible bits. For each bit position, we assigned one 20-mer sequence to encode a value of 0 and another 20-mer sequence to encode a value of 1. To aid quick hybridization, these encoding sequences were constructed from a three-letter nucleotide alphabet, one with only A, T, and C, in order to destabilize any potential secondary structures^13^. The utilized sequences were drawn from those previously used for MERFISH with additional sequences designed using approaches described previously^11^. For each barcode, the bits are concatenated with a single G separating each.

We assembled this barcode library by ligating a mixture of short, overlapping oligonucleotides. For each pair of adjacent bits, there are four unique combinations of bit values. Each corresponding sequence was synthesized as a single-stranded oligo. These oligos were then ligated to from complete, doublestranded barcodes that contain concatenated sequences of all bits with all possible bit values.

For the ligation step, all oligos were mixed and diluted so that each oligo was present at a concentration of 100 nM. The mixture was phosphorylated by incubation with T4 polynucleotide kinase (16 μL oligo mixture, 2 μL T4 ligase buffer, 2 μL PNK (NEB, M0201S)) at 37 °C for 30 minutes and ligated by adding 1 μL T4 ligase (NEB, M0202S) and incubating for 1 hour at room temperature.

To prepare a plasmid library containing these barcode sequences along with the desired promoter, we diluted the ligation product 10-fold and amplified by limited-cycle PCR on a Bio-Rad CFX96 using Phusion polymerase (NEB, M0531S0) and EvaGreen (Biotium, 3100). The PCR product was run in an agarose gel and the band of the expected length was extracted and purified (Zymo Zymoclean Gel DNA Recovery Kit, D4002). The purified product was inserted by isothermal assembly^14^ for 1 hour at 50 °C (NEB NEBuilder HiFi DNA Assembly Master Mix, E2621L) into a plasmid backbone fragment containing the colE1 origin, the ampicillin resistance gene, and other elements taken from the pZ series of plasmids^15^. The assembled plasmids were purified (Zymo DNA Clean and Concentration, D4003), eluted into 6 μL water, mixed with 10 μL of electro-competent *E. coli* on ice (NEB, C2986K), and electroporated using an Amaxa Nucleofector II. Immediately after electroporation, 1 mL SOC was added and the culture was incubated at 37 °C on a shaker for one hour. Subsequently, the SOC culture was diluted into 50 mL of LB (Teknova, L8000) supplemented with 0.1 mg/mL carbenicillin (ThermoFisher, 10177-012) and placed on the shaker at 37 °C overnight. The following day, the culture was miniprepped (Zymo Zyppy Plasmid Miniprep Kit, D4019), yielding the complete barcode library.

### Assembling protein mutant libraries

To create a library of mutant proteins, short nucleotide sequences containing regions of the protein of interest with the desired mutations were synthesized as complex oligonucleotide pools. To then create the desired mutant genes from these pools, we amplified the pool and its corresponding expression plasmid via limited cycle PCR and assembled these fragments using isothermal assembly^14^. The expression backbone was derived from the colE1 origin and the chloramphenicol resistance gene from the pZ series of plasmids^15^. Oligo pool synthesis is prone to deletions, which could lead to frameshift mutations that produce non-viable proteins. To remove these variants prior to measurement, the protein variants were translationally fused upstream to the chloramphenicol resistance protein. These constructs were electroporated into *E. coli,* as described above, and these cultures grown in the presence of chloramphenicol to select only for protein variants that did not have frame-shift mutations and which could, thus, translate component chloramphenicol resistance. These plasmids were reisolated via plasmid miniprep and the genetic variants extracted via PCR prior to combination with the barcode library.

### Merging mutation libraries with the barcode library

To merge a mutant library with the barcode library, the corresponding halves of each plasmid library were amplified by limited-cycle PCR. Of note, the forward primer for amplifying the barcode library contained 20 random nucleotides so that each assembled plasmid contained a 20-mer unique molecular identifier (UMI). Also, the protein mutant half contained the plasmid’s replication origin (colE1) while the barcode half contained the ampicillin resistance gene ensuring that only plasmids containing both halves were competent. The two halves were assembled by isothermal assemble and transfected into electrocompetent *E. coli* as described earlier. After incubating in SOC for 1 hour at 37 °C, the culture was again diluted into 50 mL and grown until it reached an OD600 of ~1. To limit the possibility that a single bacterium had taken up more than one plasmid, plasmids were extracted again from this culture, and re-electroporated into fresh *E. coli* at a defined, low concentration. This culture was then grown and rediluted to the desired number of cells assuming an OD600 of 1 corresponded to 800 million cells. The diluted culture was incubated at 37 °C overnight and the following day it was archived for future imaging experiments by diluting 1:1 in 50% glycerol (Teknova, G1796), separating into 100 μL aliquots, and stored at −80 °C. The remaining culture was mini-prepped to use as a PCR template for constructing the barcode to genotype lookup table.

### Constructing barcode to genotype lookup table

Since barcodes and mutants are assembled randomly, next generation sequencing was used to construct a look-up table from barcode to mutant. The total length of the protein mutant and the barcode exceeded the read length of the sequencing platform used (Illumina MiSeq). To circumvent this challenge, multiple fragments were extracted from each library, sequenced independently and grouped computationally using the UMI.

The mini-prepped libraries were prepared for sequencing by two sequential limited-cycle PCRs. The first PCR extracted the desired region while adding the sequencing priming regions, and the second PCR added multiplexing indices and the Illumina adapter sequences. Between PCRs, the product was purified in an agarose gel and the final product was gel purified prior to sequencing.

For each sequencing read, the corresponding barcode or mutant sequence was extracted. The reads were then grouped by common UMI and the most frequently occurring barcode and protein mutant seen for each UMI was assigned to that UMI, constructing the barcode to mutant lookup table for every variant in the library.

### Phenotype and barcode imaging

Each library was prepared for imaging by thawing the 100 μL aliquot from −80 °C to room temperature and diluting into 2 mL LB supplemented with 0.1mg/mL carbenicillin. Imaging coverslips (Bioptechs, 0420-0323-2) in 60-mm-diameter cell culture dishes were prepared by covering in 1% polyethylenimine (Sigma-Aldrich, P3143-500ML) in water for 30 minutes followed by a single wash with phosphate buffered saline (PBS). The *E. coli* culture was diluted 10-fold into PBS, poured into the culture dish, and spun at 100g for 5 minutes to adhere cells to the surface.

The sample coverslip was assembled into a Bioptech’s FCS2 flow chamber. A peristaltic pump (Gilson, MINIPULS 3) pulled liquid through the chamber while three computer-controlled valves (Hamilton, MVP and HVXM 8-5) were used to select the input fluid. The sample was imaged on a custom microscope built around a Nikon Ti-U microscope body with a Nikon CFI Plan Apo Lambda 60x oil immersion objective with 1.4 NA. Illumination was provided at 405, 488, 560, 647, and 750 nm using solid-state single-mode lasers (Coherent, Obis 405nm LX 200mW; Coherent, Genesis MX488-1000; MPB Communications, 2RU-VFL-P-2000-560-B1R, MPB Communication, 2RU-VFL-P-1500-647-B1R; and MPB Communications, 2RU-VFL-P-500-750-B1R) in addition to the overhead halogen lamp for bright field illumination. The Gaussian profile from the lasers was transformed into a top-hat profile using a refractive beam shaper (Newport, GBS-AR14). The intensity of the 488-, 560-, and 647-nm lasers was controlled by an acousto-optic tunable-filter (AOTF), the 405-nm laser was modulated by a direct digital signal, and the 750-nm laser and overhead lamp were switched by mechanical shutters. The excitation illumination was separated from the emission using a custom dichroic (Chroma, zy405/488/561/647/752RP-UF1) and emission filter (Chroma, ZET405/488/461/647-656/752m). The emission was imaged onto an Andor iXon+ 888 EMCCD camera. During acquisition, the sample was translated using a motorized XY stage (Ludl, BioPrecision2) and kept in focus using a home-built autofocus system.

Phenotype measurements were conducted immediately after cells were deposited onto the coverslip and inserted into the flow chamber. For the YFAST mutants, images were first acquired in PBS in the absence of the chromophore with 405-nm illumination to measure the mTagBFP2 fluorescence followed by an image with brightfield illumination for alignment between multiple imaging rounds. Then 10 μM HMBR or HBR in PBS was flowed over the cells and a fluorescence image was acquired with 488-nm illumination to measure YFAST and a brightfield image for alignment, followed by twenty or eighty images at 8.4Hz with constant 488-nm illumination to measure the decrease in intensity upon photobleaching. In each imaging round, images were acquired at 1,427 or 3,223 locations in the sample.

Following the phenotype measurement, the cells were fixed by incubation in a mixture of methanol and acetone at a 4:1 ratio for 30 minutes. To prevent salts from precipitating and clogging the flow system, water was flowed before and after the fixation mixture. Once fixed, the cells were washed in 2x Saline Sodium Chloride (SSC) and hybridization began.

To determine the RNA barcode expressed within each cell, MERFISH was performed using similar protocols to those described previously^11^. For each hybridization round, the sample was incubated for 30 minutes in hybridization buffer [2xSSC; 5% w/v dextran sulfate (EMD Millipore, 3730-100ML), 5% w/v ethylene carbonate (Sigma-Aldrich, E26258-500G), 0.05% w/v yeast tRNA, and 0.1% v/v Murine RNase inhibitor (NEB, M0314L)] with a mixture of readout probes labeled with either ATTO565, Cy5, or Alexa750 (Bio-Synthesis Inc) each at a concentration of 10 nM. The dyes were linked to the readout probes through a disulfide bond. Then, the hybridization buffer was replaced by an oxygen-scavenging buffer for imaging [2xSSC; 50 mM TrisHCl pH 8, 10% w/v glucose (Sigma-Aldrich, G8270), 2 mM Trolox (Sigma-Aldrich, 238813), 0.5 mg/mL glucose oxidase (Sigma-Aldrich, G2133), and 40 μg/mL catalase (Sigma-Aldrich, C100-500mg)]. Each position in the flow cell was imaged with 750-, 647-, and 560-nm illumination from longest to shortest wavelength followed by brightfield illumination before continuing to the next location. Following the imaging of all regions, the disulfide bond linking the dyes to the readout probes were cleaved by incubating the sample in 50 mM tris (2-carboxyethyl)phosphine (TCEP; Sigma-Aldrich, 646547-10X1ML) in 2xSSC for 15 minutes. The sample was then rinsed in 2xSSC and the next hybridization round started. For each round of hybridization, three readouts with spectrally discernable dyes were hybridized simultaneously. Altogether, with 14 hybridization rounds, all 42 readouts corresponding to 21 bits were measured in 40 hours. For smaller libraries, the imaging area and the number of hybridization rounds were decreased to reduce the measurement time to 22 hours.

### Image analysis

To correct for residual illumination variations across the camera, a flat field correction was performed. Every image was divided by the mean intensity image for all images with the given illumination color. Then, the images for different rounds corresponding to the same region were registered using the image acquired under bright field illumination by up-sampled cross-correlation creating a normalized image stack of all images at each position in the flow chamber. If the radial power spectral density of any given bright field image did not contain sufficient high frequency power, the image was designated as out-of-focus and all images for the corresponding region were excluded for further analysis.

To extract cell intensities, the edges of each cell were detected using the Canny edge detection algorithm on the first image acquired with 405-nm illumination. The edges that formed closed boundaries were filled in and closed regions of pixels were extracted. If a given closed pixel region had a filled area of more than 20 pixels and the ratio of the filled area to the area of the convex hull was greater than 0.9, it was classified as a cell. To increase the cell detection efficiency, the detected cells were then removed from the binary image, the image was dilated, filled, and eroded and cells were extracted again. This allowed cells where gaps exist in the detected edges to still be detected. For each cell, the mean intensity was extracted for the corresponding pixels in every image.

From the cell intensities, the phenotypes and barcodes were calculated. For each measured readout sequence, the measured intensity was normalized by subtracting the minimum and taking the median of the signal observed for that readout sequence in all cells. To determine whether a barcode contained a “1” or a “0” at each bit, the measured intensity of the “1” readout sequence and the “0” readout sequence for that bit were compared. Specifically, a threshold was selected on the ratio of these two values. If this ratio was above the threshold, the bit was called as a “1”. If it fell below, the bit was called as a “0”. Because the “1” and “0” readout sequences associated with each bit might be assigned to different fluorophores, it was necessary to optimize this threshold for each bit individually. This optimization was performed by randomly selecting 150 barcodes (a training set) from the set of known barcodes contained in the library, as identified by sequencing. An initial set of thresholds was selected and the fraction of cells matching these barcodes was determined. The threshold for each bit was then varied independently to identify the threshold set that maximizes this fraction. This optimized threshold set was then used for determining the bit values for all cells.

Once the barcode was determined for each cells, cells were grouped by barcode and the median of the various phenotype values was computed to determine the measured phenotype for the genotype corresponding to that barcode. For YFAST measurements, the normalized intensity was determined by the ratio of the YFAST fluorescence intensities under 488-nm illumination in the presence of the YFAST chromophore to the mTagBFP2 fluorescence intensities under 405-nm illumination in the absence of chromophore. To account for the non-negligible fluorescence background present in *E. coli* upon 488-nm illumination, the magnitude of the background was subtracted before calculating the fluorescence ratio. The background was estimated by calculating the median intensity of all cells upon 488-nm illumination predicted to contain a non-fluorescent YFAST mutant. Cells, grouped by barcode, were assigned to the dark population if the Pearson correlation coefficient between the 488-nm intensity and the 405-nm intensity for the grouped cells fell below the arbitrary threshold of 0.2. When the two intensities are uncorrelated, it suggests the number of YFAST proteins in the cells associated with that barcode does not affect the brightness of the cell and hence the YFAST should be dark. To determine the relative amplitude of the fast decay component of the photobleaching measurement of each YFAST mutant, we fit the background subtracted photobleaching curve to the sum of two exponentials, with one of the decay rates set to the fast decay rate determined from the original YFAST. To determine the fast decay rate of the original YFAST, its background subtracted photobleaching curve was fit to the sum of two exponentials with both decay rates as adjustable parameters and the faster of the two decay rates was selected.

